# Increased hippocampal epigenetic age in the Ts65Dn Mouse Model of Down Syndrome

**DOI:** 10.1101/2023.09.25.559272

**Authors:** Francesco Ravaioli, Fiorenza Stagni, Sandra Guidi, Chiara Pirazzini, Paolo Garagnani, Alessandro Silvani, Giovanna Zoccoli, Renata Bartesaghi, Maria Giulia Bacalini

## Abstract

Down syndrome (DS) is a segmental progeroid genetic disorder associated to multi-systemic precocious ageing phenotypes, which are particularly evident at the immune and nervous systems. Accordingly, people with DS show an increased biological age as measured by epigenetic clocks. Ts65Dn trisomic mouse, which harbors extra-numerary copies of Hsa21-syntenic regions, was shown to recapitulate several progeroid features of DS, but no biomarkers of age have been applied to it so far. Here we used a mouse specific epigenetic clock to measure epigenetic age of hippocampi from Ts65Dn and euploid mice at 20 weeks. Ts65Dn mice showed an increased hippocampal epigenetic age respect to controls, and the observed changes in DNA methylation partially recapitulated those observed in hippocampi from people with DS. Collectively, our results support the use of the Ts65Dn model to decipher the molecular mechanisms underlying the progeroid DS phenotypes.

## Increased hippocampal epigenetic age in the Ts65Dn Mouse Model of Down Syndrome

Down Syndrome (DS) is a common genetic disorder caused by complete or segmental triplication of chromosome 21 (Hsa21) and is the most frequent genetic cause of intellectual disability. DS is considered a segmental progeroid syndrome, characterized by a precocious aging-like deterioration which is particularly evident at the immune system and brain level. This view, originally proposed by George Martin on the basis of the analysis of DS phenotypic traits (Martin 1978), has been further refined in the last two decades through physiological and molecular analyses that explored similarities and differences between the pillars of aging and alterations occurring in DS (Franceschi et al. 2019; Chen et al. 2021; Zigman 2013). In this framework, several biomarkers of age have been explored in people with DS, including those based on telomere length (Gimeno et al. 2014; Holmes et al. 2006), magnetic resonance neuroimaging (brain-age) (Cole et al. 2017), serum proteins glycosylation (GlycoAge) (Borelli et al. 2015) and DNA methylation (DNAm) (epigenetic clocks) (Horvath, Garagnani, et al. 2015). These studies concordantly suggest that people with DS are older than their chronological age.

Murine models are largely employed in the study of ageing and age-related disease (Palliyaguru et al. 2021) and mouse epigenetic biomarkers of age have been developed (Coninx et al. 2020; Lu et al. 2023; Han et al. 2018; Zhou et al. 2022; Wang & Lemos 2019). So far, however, these epigenetic clocks have been applied to a limited extent and, to the best of our knowledge, no data are available for mouse models of DS.

The Ts65Dn mouse strain is the most common model for the study of DS. These mice are segmentally trisomic for a region of chromosome 16 that is homologous to part of Hsa21. Ts65Dn mice were shown to recapitulate a wide range of DS-specific behavioral, physiological and neuroanatomical features such as reduced brain size, neuronal density (Lorenzi & Reeves 2006; Baxter 2000; Insausti et al. 1998; Stagni et al. 2018), altered neuronal function (Kleschevnikov et al. 2004; Siarey et al. 1997) and altered dendrite architecture in hippocampal regions (Uguagliati et al. 2022) as well as spatial learning and memory deficits. The Ts65Dn mouse, in addition, shares with the DS human condition multi-systemic premature aging associated with early alterations in mitochondrial function, DNA damage response, proteostasis and early neurodegeneration (Cisterna et al. 2020; Vacano et al. 2012; Puente-Bedia et al. 2022; Mollo et al. 2020; Holtzman et al. 1996; Kirstein et al. 2022).

In this study, we aimed to evaluate hippocampal epigenetic age in the Ts65Dn model. We used a hippocampus-specific mouse epigenetic clock developed by Zymo Research (referred as DNAge®) which is based on deep bisulfite sequencing of 300 target regions containing 2045 CpG sites. Using this clock, Coninx and colleagues previously reported an increase in epigenetic age in the triple transgenic AD mouse model (Coninx et al. 2020).

We applied the DNAge® clock to 6 Ts65Dn mice (age: 20 weeks; 5 males and 1 female) and 7 euploid mice (age: 20 weeks, 6 males and 1 female) (Supplementary information). As shown in Figure 1A, the hippocampal epigenetic clock model tended to underestimate the age of euploid mice (mean epigenetic age: 4.7 weeks instead of 20), an effect that is in line with the original publication (Coninx et al. 2020). With respect to the estimated epigenetic age of euploid controls, Ts65Dn mice were significantly epigenetically older (Mann-Whitney test p-value=0.0047). This result indicates for the first time that the Ts65Dn murine model mimics the increase in epigenetic age previously described in the brain and blood of subjects with DS (Horvath, Garagnani, et al. 2015; Do et al. 2017). Ts65Dn mice also showed a higher variance compared to euploid mice, although not reaching statistical significance (F-test p-value = 0.1231). This trend in higher variance can be related to the progeroid phenotype, as an increase in epigenetic variability has been described during aging (BIOS consortium et al. 2016), although we cannot exclude that it is the result of the phenotypic drift observed in the Ts65Dn model (Shaw et al. 2020).

**Figure 1.**
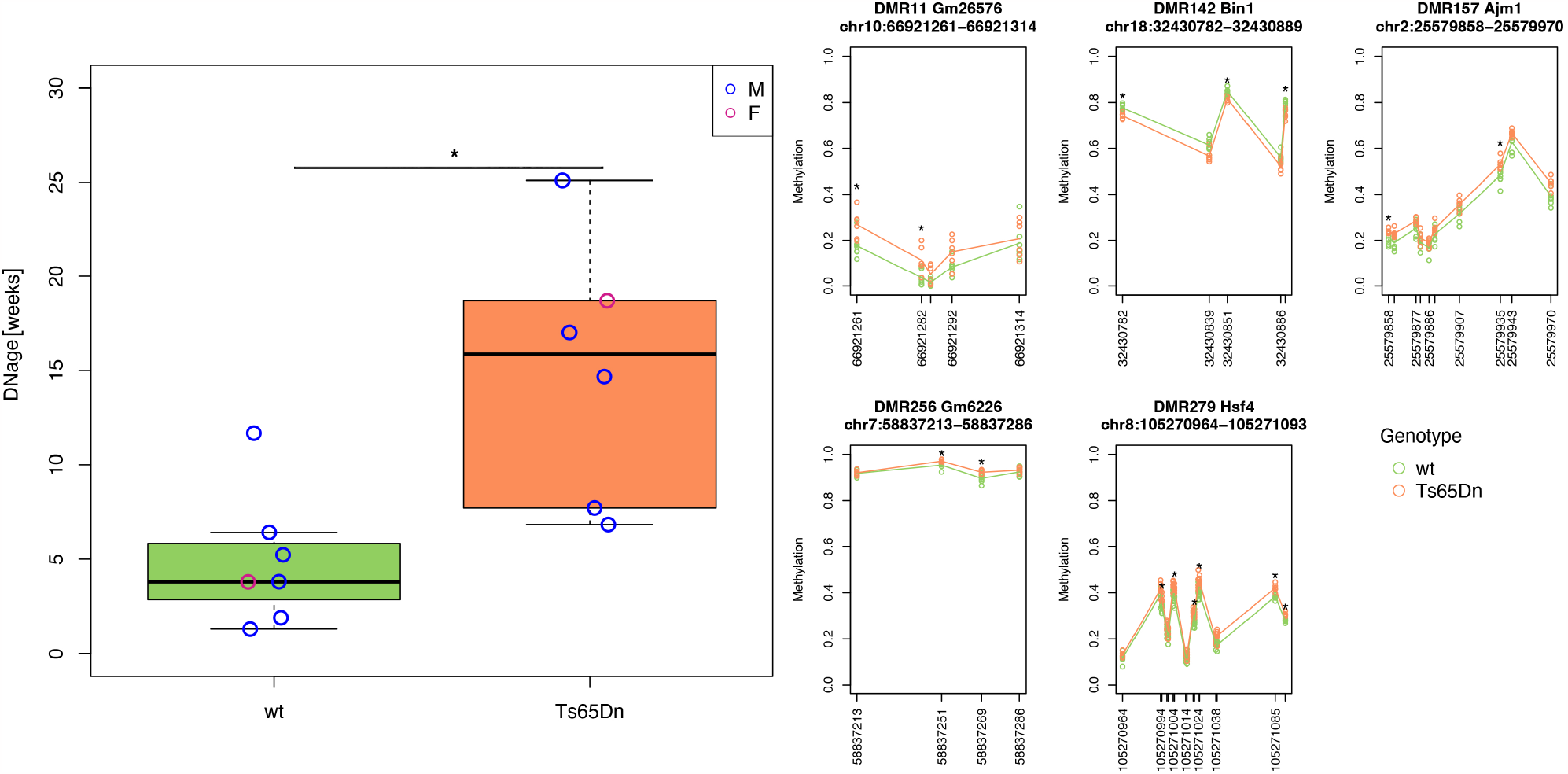
Increased epigenetic age of hippocampi in Ts65Dn mice. Boxplots showing epigenetic age predicted using ZymoResearch DNAge® predictor algorithm in Ts65Dn and wild-type euploid (wt) mice. Blue and red circles indicate data from male and female mice, respectively. *: p < 0.01, Mann-Whitney test). B) Lineplots of DNA methylation profiles in Ts65Dn mouse hippocampi for differentially methylated regions with at least two significant CpG sites. *: p < 0.01, Mann-Whitney test).

To get further insights into the DNAm differences that contribute to the increased epigenetic age of Ts65Dn mice, we applied the Mann-Whitney test to each CpG site that makes the epigenetic clock. We found 27 differentially methylated CpGs (nominal p-value<0.01), of which 21 were hypermethylated and 6 hypomethylated in Ts65Dn compared to euploid mice (Supplementary Table 1). Furthermore, we identified 5 genomic regions that contained at least two CpG sites having a nominal p-value<0.01 (Figure 1B). These differentially methylated regions (DMRs) annotated to *Bin1, Ajm1, Hsf4, Gm2662* and *Gm26576* genes. *Bin1* is a ubiquitously expressed gene which is known to modulate tau processing as well as to be involved in vesicle trafficking, inflammation and apoptosis (GERAD consortium et al. 2013; Thinakaran & Koo 2008; Galderisi et al. 1999). *Ajm1* seems to be involved in cell-to-cell organization, while *Hsf4* is a transcription factor known to act upstream of several processes, including DNA damage repair (Cui et al. 2012), and is central in the development of the eye (Fujimoto et al. 2004). *Gm2662* and *Gm26576* functions are not known.

Previous studies using whole-genome bisulfite sequencing (WGBS) on whole cerebral hemispheres of newborn Dp(16)1Yey and Dp(10)1Yey mice highlighted similarities in DNAm profiles between trisomic humans and mice (Mendioroz et al. 2015). We therefore performed a cross-species analysis to check whether the DNAge® CpG sites differentially methylated in Ts65Dn mice showed altered methylation in hippocampi from subjects with DS. We searched Gene Expression Omnibus (GEO) repository and found a small dataset (GSE63347) containing DNAm data from hippocampi from 2 subjects with DS (age:42-57 y.o., 2 males) and 7 euploid controls (age:38-64, 2 males and 5 females), generated by the *Illumina Infinium HumanMethylation450K* microarray (Horvath, Garagnani, et al. 2015). Using UCSC *lift-over* tool (Hinrichs 2006), the genomic coordinates of 15 out of the 27 CpG sites identified above were lifted from mouse (mm10 genome assembly) to human (hg19 genome assembly). We analyzed all the human microarray probes mapping between 250bp upstream and 250bp downstream of the 15 lifted CpG sites (a total of 19 probes). No probe was significantly differentially methylated between subjects with DS and controls (Mann-Whitney test p-value>0.05), possibly due also to the small sample size of the dataset; however, we found a CpG probe (cg04235075) mapping within *HSF4* gene which showed a trend towards hypermethylation in DS (Figure 2A), concordantly with what observed in mouse. Interestingly, DNAm of this probe was also found to be positively associated with age in healthy human hippocampi, as resulting from the analysis of GSE129428 (age 34-78, 13 females and 19 males) (Fries et al. 2020) and GSE64509 (age 38-114, 17 females and 8 males) (Horvath, Mah, et al. 2015) datasets (Figure 2B and Figure 2C). This concordance in DNAm changes observed in trisomic mice and humans as well as in human aging is of interest, as it suggests the presence of cross-species conserved epigenetic mechanisms that can contribute to the progeroid phenotype of DS.

**Figure 2.**
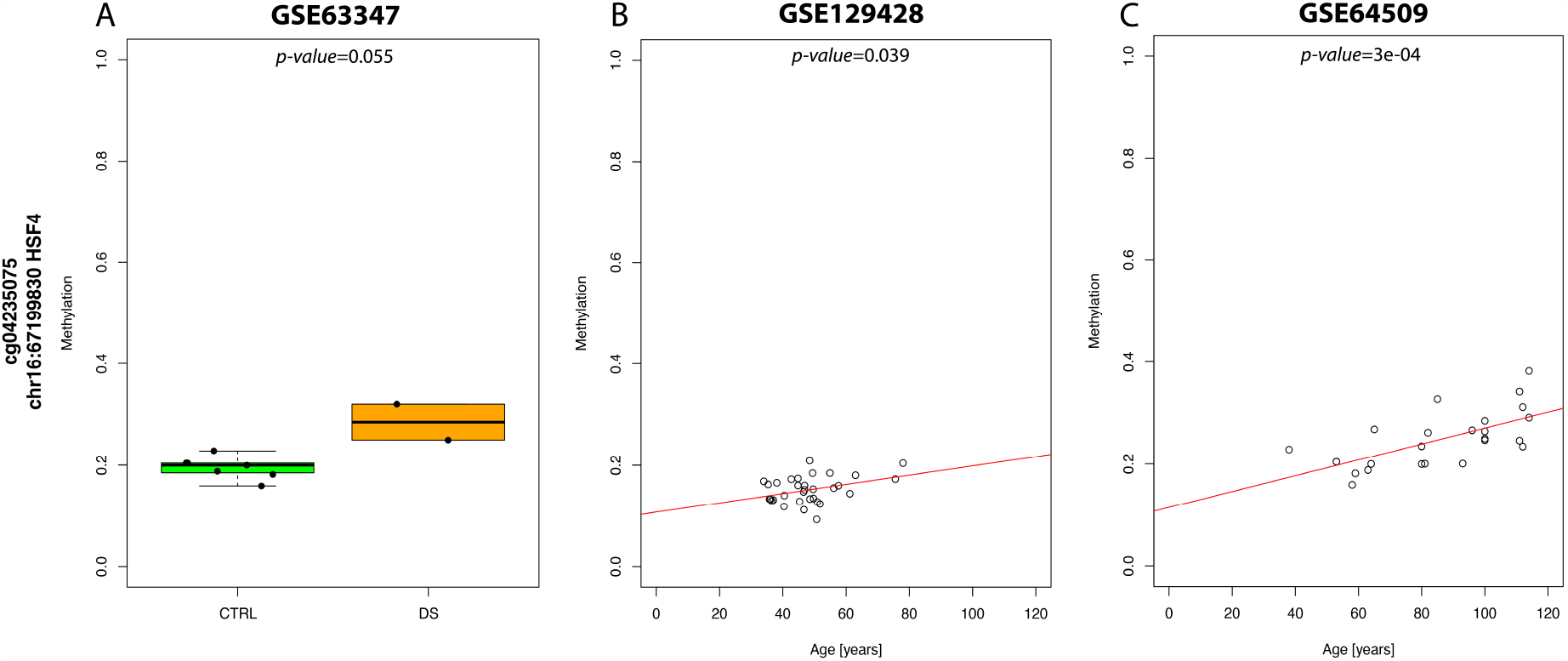
DNAm profiles of *HSF4* CpGs in human hippocampi. A) DNAm profiles of Illumina Infinium 450k probes cg03140421 and cg04235075, mapping within human Hsf4 orthologous region, in hippocampus from subjects with DS (GSE63347) (*p-value* calculated with Mann-Whitney test). B, C) DNAm profiles of Illumina Infinium 450k probes cg03140421 and cg04235075 in hippocampus from subjects without overt pathologies at different ages (GSE129428, GSE64509) (*p-values* calculated with linear model).

Collectively, our results show an increased hippocampal epigenetic age in the Ts65Dn mouse model. This is fully in line with its progeroid phenotypes (see above) and the notion that epigenetic alterations represent a pillar of aging (Mendioroz et al. 2015). The analysis of a small DNAm dataset of hippocampi from DS subjects revealed that some DNAm changes found in Ts65Dn mice were also present in humans. Although larger samples are needed, our study supports the use of the Ts65Dn model to decipher the molecular mechanisms underlying the progeroid DS phenotype. Finally, our study supports the use of the DNAge® clock and, possibly, other recently developed mouse epigenetic clocks (Lu et al. 2023; Zhou et al. 2022), as biomarkers of biological age. Such tools might be exploited to monitor the impact of disease and disease-modifying interventions in DS. It is worth to note that the DNAge® epigenetic clock is based on deep bisulfite sequencing of few genomic regions and therefore it is not informative of genome-wide epigenetic remodeling occurring in Ts65Dn mice. The recent release of *Illumina Infinium Mouse Methylation* microarray will allow investigation of genome wide DNAm profiles of DS models in a cost-effective manner and to get further insights into the epigenetic basis of the progeroid phenotype of DS.

## Abbreviations

DS–: Down Syndrome
Hsa21–: Human Chromosome 21
DNAm–: DNA methylation

## Authors’ Contributions

Maria Giulia Bacalini, Renata Bartesaghi and Fiorenza Stagni conceptualized and designed the study; Fiorenza Stagni and Sandra Guidi provided tissue samples; Francesco Ravaioli and Maria Giulia Bacalini performed experimental procedures and statistical analysis; Maria Giulia Bacalini, and Francesco Ravaioli drafted the manuscript; Maria Giulia Bacalini, Renata Bartesaghi, Paolo Garagnani, Sandra Guidi, Chiara Pirazzini, Alessandro Silvani; Fiorenza Stagni and Giovanna Zoccoli contributed to the interpretation of results and the review of the manuscript. All authors approved the final version of the manuscript.

## References

Baxter LL (2000) Discovery and genetic localization of Down syndrome cerebellar phenotypes using the Ts65Dn mouse. Human Molecular Genetics 9, 195–202.DOI: 10.1093/hmg/9.2.195

BIOS consortium, Slieker RC, van Iterson M, Luijk R, Beekman M, Zhernakova DV, Moed MH, Mei H, van Galen M, Deelen P, Bonder MJ, Zhernakova A, Uitterlinden AG, Tigchelaar EF, Stehouwer CDA, Schalkwijk CG, van der Kallen CJH, Hofman A, van Heemst D, de Geus EJ, van Dongen J, Deelen J, van den Berg LH, van Meurs J, Jansen R, ‘t Hoen PAC, Franke L, Wijmenga C, Veldink JH, Swertz MA, van Greevenbroek MMJ, van Duijn CM, Boomsma DI, Slagboom PE & Heijmans BT (2016) Age-related accrual of methylomic variability is linked to fundamental ageing mechanisms. Genome Biology 17.DOI: 10.1186/s13059-016-1053-6 Available at: http://genomebiology.biomedcentral.com/articles/10.1186/s13059-016-1053-6 [Accessed May 6, 2017].

Borelli V, Vanhooren V, Lonardi E, Reiding KR, Capri M, Libert C, Garagnani P, Salvioli S, Franceschi C & Wuhrer M (2015) Plasma N-Glycome Signature of Down Syndrome. J Proteome Res 14, 4232–4245.DOI: 10.1021/acs.jproteome.5b00356

Chen X-Q, Xing Z, Chen Q-D, Salvi RJ, Zhang X, Tycko B, Mobley WC & Yu YE (2021) Mechanistic Analysis of Age-Related Clinical Manifestations in Down Syndrome. Front. Aging Neurosci. 13, 700280.DOI: 10.3389/fnagi.2021.700280

Cisterna B, Sobolev AP, Costanzo M, Malatesta M & Zancanaro C (2020) Combined Microscopic and Metabolomic Approach to Characterize the Skeletal Muscle Fiber of the Ts65Dn Mouse, A Model of Down Syndrome. Microsc Microanal 26, 1014–1023.DOI: 10.1017/S143192762002437X

Cole JH, Annus T, Wilson LR, Remtulla R, Hong YT, Fryer TD, Acosta-Cabronero J, Cardenas-Blanco A, Smith R, Menon DK, Zaman SH, Nestor PJ & Holland AJ (2017) Brain-predicted age in Down syndrome is associated with beta amyloid deposition and cognitive decline. Neurobiology of Aging 56, 41–49.DOI: 10.1016/j.neurobiolaging.2017.04.006

Coninx E, Chew YC, Yang X, Guo W, Coolkens A, Baatout S, Moons L, Verslegers M & Quintens R (2020) Hippocampal and cortical tissue-specific epigenetic clocks indicate an increased epigenetic age in a mouse model for Alzheimer’s disease. aging 12, 20817–20834.DOI: 10.18632/aging.104056

Cui X, Zhang J, Du R, Wang L, Archacki S, Zhang Y, Yuan M, Ke T, Li H, Li D, Li C, Li DW-C, Tang Z, Yin Z & Liu M (2012) HSF4 is involved in DNA damage repair through regulation of Rad51. Biochimica et Biophysica Acta (BBA) - Molecular Basis of Disease 1822, 1308–1315.DOI: 10.1016/j.bbadis.2012.05.005

Do C, Xing Z, Yu YE & Tycko B (2017) Trans-acting epigenetic effects of chromosomal aneuploidies: lessons from Down syndrome and mouse models. Epigenomics 9, 189–207.DOI: 10.2217/epi-2016-0138

Franceschi C, Garagnani P, Gensous N, Bacalini MG, Conte M & Salvioli S (2019) Accelerated bio-cognitive aging in Down syndrome: State of the art and possible deceleration strategies. Aging Cell, e12903.DOI: 10.1111/acel.12903

Fries GR, Bauer IE, Scaini G, Valvassori SS, Walss-Bass C, Soares JC & Quevedo J (2020) Accelerated hippocampal biological aging in bipolar disorder. Bipolar Disorders 22, 498–507.DOI: 10.1111/bdi.12876

Fujimoto M, Izu H, Seki K, Fukuda K, Nishida T, Yamada S, Kato K, Yonemura S, Inouye S & Nakai A (2004) HSF4 is required for normal cell growth and differentiation during mouse lens development. EMBO J 23, 4297–4306.DOI: 10.1038/sj.emboj.7600435

Galderisi U, Di Bernardo G, Cipollaro M, Jori FP, Piegari E, Cascino A, Peluso G & Melone MA (1999) Induction of apoptosis and differentiation in neuroblastoma and astrocytoma cells by the overexpression of Bin1, a novel Myc interacting protein. J Cell Biochem 74, 313–322.

GERAD consortium, Chapuis J, Hansmannel F, Gistelinck M, Mounier A, Van Cauwenberghe C, Kolen KV, Geller F, Sottejeau Y, Harold D, Dourlen P, Grenier-Boley B, Kamatani Y, Delepine B, Demiautte F, Zelenika D, Zommer N, Hamdane M, Bellenguez C, Dartigues J-F, Hauw J-J, Letronne F, Ayral A-M, Sleegers K, Schellens A, Broeck LV, Engelborghs S, De Deyn PP, Vandenberghe R, O’Donovan M, Owen M, Epelbaum J, Mercken M, Karran E, Bantscheff M, Drewes G, Joberty G, Campion D, Octave J-N, Berr C, Lathrop M, Callaerts P, Mann D, Williams J, Buée L, Dewachter I, Van Broeckhoven C, Amouyel P, Moechars D, Dermaut B & Lambert J-C (2013) Increased expression of BIN1 mediates Alzheimer genetic risk by modulating tau pathology. Mol Psychiatry 18, 1225–1234.DOI: 10.1038/mp.2013.1

Gimeno A, García-Giménez JL, Audí L, Toran N, Andaluz P, Dasí F, Viña J & Pallardó FV (2014) Decreased cell proliferation and higher oxidative stress in fibroblasts from Down Syndrome fetuses. Preliminary study. Biochimica et Biophysica Acta (BBA) - Molecular Basis of Disease 1842, 116–125.DOI: 10.1016/j.bbadis.2013.10.014

Han Y, Eipel M, Franzen J, Sakk V, Dethmers-Ausema B, Yndriago L, Izeta A, de Haan G, Geiger H & Wagner W (2018) Epigenetic age-predictor for mice based on three CpG sites. eLife 7, e37462.DOI: 10.7554/eLife.37462

Hinrichs AS (2006) The UCSC Genome Browser Database: update 2006. Nucleic Acids Research 34, D590–D598.DOI: 10.1093/nar/gkj144

Holmes DK, Bates N, Murray M, Ladusans EJ, Morabito A, Bolton-Maggs PHB, Johnston TA, Walkenshaw S, Wynn RF & Bellantuono I (2006) Hematopoietic progenitor cell deficiency in fetuses and children affected by Down’s syndrome. Experimental Hematology 34, 1611–1615.DOI: 10.1016/j.exphem.2006.10.013

Holtzman DM, Santucci D, Kilbridge J, Chua-Couzens J, Fontana DJ, Daniels SE, Johnson RM, Chen K, Sun Y, Carlson E, Alleva E, Epstein CJ & Mobley WC (1996) Developmental abnormalities and age-related neurodegeneration in a mouse model of Down syndrome. Proc. Natl. Acad. Sci. U.S.A. 93, 13333–13338.DOI: 10.1073/pnas.93.23.13333

Horvath S, Garagnani P, Bacalini MG, Pirazzini C, Salvioli S, Gentilini D, Di Blasio AM, Giuliani C, Tung S, Vinters HV & Franceschi C (2015) Accelerated epigenetic aging in Down syndrome. Aging Cell 14, 491–495.DOI: 10.1111/acel.12325

Horvath S, Mah V, Lu AT, Woo JS, Choi O-W, Jasinska AJ, Riancho JA, Tung S, Coles NS, Braun J, Vinters HV & Coles LS (2015) The cerebellum ages slowly according to the epigenetic clock. Aging (Albany NY) 7, 294–306.DOI: 10.18632/aging.100742

Insausti AM, Megias M, Crespo D, Cruz-Orive LM, Dierssen M, Vallina TF, Insausti R & Flórez J (1998) Hippocampal volume and neuronal number in Ts65Dn mice: a murine model of down syndrome. Neuroscience Letters 253, 175–178.DOI: 10.1016/S0304-3940(98)00641-7

Kirstein M, Cambrils A, Segarra A, Melero A & Varea E (2022) Cholinergic Senescence in the Ts65Dn Mouse Model for Down Syndrome. Neurochem Res 47, 3076–3092.DOI: 10.1007/s11064-022-03659-0

Kleschevnikov AM, Belichenko PV, Villar AJ, Epstein CJ, Malenka RC & Mobley WC (2004) Hippocampal Long-Term Potentiation Suppressed by Increased Inhibition in the Ts65Dn Mouse, a Genetic Model of Down Syndrome. J. Neurosci. 24, 8153–8160.DOI: 10.1523/JNEUROSCI.1766-04.2004

Lorenzi HA & Reeves RH (2006) Hippocampal hypocellularity in the Ts65Dn mouse originates early in development. Brain Research 1104, 153–159.DOI: 10.1016/j.brainres.2006.05.022

Lu AT, Fei Z, Haghani A, Robeck TR, Zoller JA, Li CZ, Lowe R, Yan Q, Zhang J, Vu H, Ablaeva J, Acosta-Rodriguez VA, Adams DM, Almunia J, Aloysius A, Ardehali R, Arneson A, Baker CS, Banks G, Belov K, Bennett NC, Black P, Blumstein DT, Bors EK, Breeze CE, Brooke RT, Brown JL, Carter GG, Caulton A, Cavin JM, Chakrabarti L, Chatzistamou I, Chen H, Cheng K, Chiavellini P, Choi OW, Clarke SM, Cooper LN, Cossette ML, Day J, DeYoung J, DiRocco S, Dold C, Ehmke EE, Emmons CK, Emmrich S, Erbay E, Erlacher-Reid C, Faulkes CG, Ferguson SH, Finno CJ, Flower JE, Gaillard JM, Garde E, Gerber L, Gladyshev VN, Gorbunova V, Goya RG, Grant MJ, Green CB, Hales EN, Hanson MB, Hart DW, Haulena M, Herrick K, Hogan AN, Hogg CJ, Hore TA, Huang T, Izpisua Belmonte JC, Jasinska AJ, Jones G, Jourdain E, Kashpur O, Katcher H, Katsumata E, Kaza V, Kiaris H, Kobor MS, Kordowitzki P, Koski WR, Krützen M, Kwon SB, Larison B, Lee SG, Lehmann M, Lemaitre JF, Levine AJ, Li C, Li X, Lim AR, Lin DTS, Lindemann DM, Little TJ, Macoretta N, Maddox D, Matkin CO, Mattison JA, McClure M, Mergl J, Meudt JJ, Montano GA, Mozhui K, Munshi-South J, Naderi A, Nagy M, Narayan P, Nathanielsz PW, Nguyen NB, Niehrs C, O’Brien JK, O’Tierney Ginn P, Odom DT, Ophir AG, Osborn S, Ostrander EA, Parsons KM, Paul KC, Pellegrini M, Peters KJ, Pedersen AB, Petersen JL, Pietersen DW, Pinho GM, Plassais J, Poganik JR, Prado NA, Reddy P, Rey B, Ritz BR, Robbins J, Rodriguez M, Russell J, Rydkina E, Sailer LL, Salmon AB, Sanghavi A, Schachtschneider KM, Schmitt D, Schmitt T, Schomacher L, Schook LB, Sears KE, Seifert AW, Seluanov A, Shafer ABA, Shanmuganayagam D, Shindyapina AV, Simmons M, Singh K, Sinha I, Slone J, Snell RG, Soltanmaohammadi E, Spangler ML, Spriggs MC, Staggs L, Stedman N, Steinman KJ, Stewart DT, Sugrue VJ, Szladovits B, Takahashi JS, Takasugi M, Teeling EC, Thompson MJ, Van Bonn B, Vernes SC, Villar D, Vinters HV, Wallingford MC, Wang N, Wayne RK, Wilkinson GS, Williams CK, Williams RW, Yang XW, Yao M, Young BG, Zhang B, Zhang Z, Zhao P, Zhao Y, Zhou W, Zimmermann J, Ernst J, Raj K & Horvath S (2023) Universal DNA methylation age across mammalian tissues. Nat Aging.DOI: 10.1038/s43587-023-00462-6 Available at: https://www.nature.com/articles/s43587-023-00462-6 [Accessed September 5, 2023].

Martin GM (1978) Genetic syndromes in man with potential relevance to the pathobiology of aging. Birth Defects Orig Artic Ser 14, 5–39.

Mendioroz M, Do C, Jiang X, Liu C, Darbary HK, Lang CF, Lin J, Thomas A, Abu-Amero S, Stanier P, Temkin A, Yale A, Liu M-M, Li Y, Salas M, Kerkel K, Capone G, Silverman W, Yu YE, Moore G, Wegiel J & Tycko B (2015) Trans effects of chromosome aneuploidies on DNA methylation patterns in human Down syndrome and mouse models. Genome Biol 16, 263.DOI: 10.1186/s13059-015-0827-6

Mollo N, Cicatiello R, Aurilia M, Scognamiglio R, Genesio R, Charalambous M, Paladino S, Conti A, Nitsch L & Izzo A (2020) Targeting Mitochondrial Network Architecture in Down Syndrome and Aging. IJMS 21, 3134.DOI: 10.3390/ijms21093134

Palliyaguru DL, Vieira Ligo Teixeira C, Duregon E, Di Germanio C, Alfaras I, Mitchell SJ, Navas-Enamorado I, Shiroma EJ, Studenski S, Bernier M, Camandola S, Price NL, Ferrucci L & De Cabo R (2021) Study of Longitudinal Aging in Mice: Presentation of Experimental Techniques R. M. Anderson, ed. The Journals of Gerontology: Series A 76, 552–560.DOI: 10.1093/gerona/glaa285

Puente-Bedia A, Berciano MT, Martínez-Cué C, Lafarga M & Rueda N (2022) Oxidative-Stress-Associated Proteostasis Disturbances and Increased DNA Damage in the Hippocampal Granule Cells of the Ts65Dn Model of Down Syndrome. Antioxidants 11, 2438.DOI: 10.3390/antiox11122438

Shaw PR, Klein JA, Aziz NM & Haydar TF (2020) Longitudinal neuroanatomical and behavioral analyses show phenotypic drift and variability in the Ts65Dn mouse model of Down syndrome. Disease Models & Mechanisms, dmm.046243.DOI: 10.1242/dmm.046243

Siarey RJ, Stoll J, Rapoport SI & Galdzicki Z (1997) Altered long-term potentiation in the young and old Ts65Dn mouse, a model for down syndrome. Neuropharmacology 36, 1549–1554.DOI: 10.1016/S0028-3908(97)00157-3

Stagni F, Giacomini A, Emili M, Guidi S & Bartesaghi R (2018) Neurogenesis impairment: An early developmental defect in Down syndrome. Free Radical Biology and Medicine 114, 15–32.DOI: 10.1016/j.freeradbiomed.2017.07.026

Thinakaran G & Koo EH (2008) Amyloid Precursor Protein Trafficking, Processing, and Function. Journal of Biological Chemistry 283, 29615–29619.DOI: 10.1074/jbc.R800019200

Uguagliati B, Stagni F, Emili M, Giacomini A, Russo C, Guidi S & Bartesaghi R (2022) Early Appearance of Dendritic Alterations in Neocortical Pyramidal Neurons of the Ts65Dn Model of Down Syndrome. Dev Neurosci 44, 23–38.DOI: 10.1159/000520925

Vacano GN, Duval N & Patterson D (2012) The Use of Mouse Models for Understanding the Biology of Down Syndrome and Aging. Current Gerontology and Geriatrics Research 2012, 1–20.DOI: 10.1155/2012/717315

Wang M & Lemos B (2019) Ribosomal DNA harbors an evolutionarily conserved clock of biological aging. Genome Res. 29, 325–333.DOI: 10.1101/gr.241745.118

Zhou W, Hinoue T, Barnes B, Mitchell O, Iqbal W, Lee SM, Foy KK, Lee K-H, Moyer EJ, VanderArk A, Koeman JM, Ding W, Kalkat M, Spix NJ, Eagleson B, Pospisilik JA, Szabó PE, Bartolomei MS, Vander Schaaf NA, Kang L, Wiseman AK, Jones PA, Krawczyk CM, Adams M, Porecha R, Chen BH, Shen H & Laird PW (2022) DNA methylation dynamics and dysregulation delineated by high-throughput profiling in the mouse. Cell Genomics 2, 100144.DOI: 10.1016/j.xgen.2022.100144

Zigman WB (2013) Atypical aging in down syndrome: Atypical Aging in Down Syndrome. Dev. Disabil. Res. Rev. 18, 51–67.DOI: 10.1002/ddrr.1128

