## Supplementary Information for "Increased hippocampal epigenetic age in the Ts65Dn Mouse Model of Down Syndrome"

***Sample Collection***

Ts65Dn and euploid mice were generated by crossing B6EiC3Sn a/A-Ts(17^16) 65Dn females (JAX line 1924) with C57BL/6JEiJ × C3H/HeSnJ (B6EiC3Sn) F1 hybrid males (JAX line 1875; euploid males) that were supplied by Jackson Laboratories (Bar Harbor, ME, USA). Ts65Dn (nMales=5, nFemales=1) and euploid (nMales=6, nFemales=1) littermates aged 20 weeks were anesthetized, their hippocampi were quickly collected and immediately snap-frozen in liquid nitrogen. DNA was isolated using AllPrep DNA/RNA/miRNA Universal Kit (QIAGEN), quantified by the QuBit DNA Broad Range kit (Thermo Fisher) and stabilized using DNA/RNA Shield reagent (1:3 ratio) for downstream processing.

***Bisulfite Targeted Sequencing***

DNA bisulfite conversion, library preparation, sequencing and data analysis for DNAge® clock were performed by the service provider (Zymo Research). Genomic DNA recovered from DNA/RNA shield solution was bisulfite-converted using EZ DNA Methylation-Lightning Kit (Zymo Research, Irvine, CA). Sequencing libraries were prepared according to the Simplified Whole-panel Amplification Reaction Method (SWARM®) which allows for enrichment of age-associated genomic loci counting more than 2000 CpG sites. Sequencing was run on an Illumina Novaseq platform for at least 1000X sequencing depth per CpG position. Sequences were identified using an Illumina base-caller software and then aligned to the reference genome (GRCm38/mm10) using Bismark. Methylation levels for each assayed cytosine were calculated as the ratio between the number of reads reporting a C and the number of reads reporting either a C or a T (beta value).

***Analysis of DNAm data***

All analyses were performed in R (v 4.2.3). Differences in DNAge® predicted epigenetic age between Ts65Dn and euploid mice were evaluated using the Mann-Whitney test. The comparison between DNAge® predicted epigenetic age variance between Ts65Dn and euploid mice was assessed using the F-test. To assess DNAm differences between Ts65Dn and euploid mice at single CpG level, DNAm beta values obtained from Zymo Research sequencing service were converted to M-values using *logit2* function of *minfi* R package. Then, differences in DNAm levels of each CpG site making up the DNAge epigenetic clock (n=2040) were evaluated using the Mann-Whitney test. Differentially methylated regions (DMRs) were defined as regions containing at least two significant CpG sites less than 200 bp apart from each other.

***Integration with human hippocampal DNAm data***

DNAm data from human hippocampi samples were downloaded from Gene Expression Omnibus (GEO) repository. To convert genomic coordinates of DNAge® epigenetic clock from mouse (mm10) to human (hg19), the *liftOver* UCSC tool was used. Since most of successfully converted murine CpG sites did not overlap with any Illumina Infinium 450k CpG probe in humans, we considered all human microarray probes mapping within 250bp (upstream or downstream) of converted murine CpG site. The GSE63347 dataset contains Infinium 450K DNAm data from hippocampi from 2 subjects with DS (age:42-57 years, 2 males) and 7 euploid controls (age:38-64, 2 males and 5 females). DNAm beta values were converted to M-values using *logit2* function of *minfi* R package and compared between DS and euploid controls using the Mann- Whitney test. GSE129428 and GSE64509 contain Illumina Infinium 450K DNAm data respectively from 25 (age range: 34-78 years) and 32 (age range: 38-114 years) hippocampi from subjects without any overt pathology. The association with age was calculated by fitting a linear model to each CpG probe (converted to M values as above) using *limma* R package.

***Supplementary Table 1***

| Chrom | Position | Gene | DMR | Illumina.ProbeID | Mann-Whitney | Delta_Beta |
|---|---|---|---|---|---|---|
| chr2 | 25579858 | *Ajm1* | DMR157 | NA | 0.004662 | 0.03481325 |
| chr2 | 25579935 | *Ajm1* | DMR157 | NA | 0.00815851 | 0.04655926 |
| chr3 | 53477651 | *Proser1* | DMR178 | NA | 0.00815851 | -0.0628105 |
| chr4 | 114666147 | *NA* | DMR197 | NA | 0.00815851 | 0.01935955 |
| chr4 | 115978859 | *Faah* | DMR198 | cg41311474 | 0.004662 | 0.04512902 |
| chr5 | 111418977 | *Mn1* | DMR217 | NA | 0.00815851 | 0.03648862 |
| chr7 | 44959987 | *Cpt1c* | DMR254 | NA | 0.00815851 | -0.0662964 |
| chr7 | 58837251 | *Gm6226* | DMR256 | NA | 0.002331 | 0.01700904 |
| chr7 | 58837269 | *Gm6226* | DMR256 | NA | 0.00815851 | 0.02677204 |
| chr7 | 80876214 | *Zscan2* | DMR259 | NA | 0.00815851 | 0.06676135 |
| chr7 | 141269419 | *Cdhr5* | DMR267 | NA | 0.002331 | 0.09361614 |
| chr8 | 88664327 | *Nod2* | DMR275 | NA | 0.00815851 | 0.02826343 |
| chr8 | 105270995 | *Hsf4* | DMR279 | cg46095458 | 0.0011655 | 0.03989681 |
| chr8 | 105271005 | *Hsf4* | DMR279 | NA | 0.0011655 | 0.04671537 |
| chr8 | 105271021 | *Hsf4* | DMR279 | NA | 0.0011655 | 0.0376626 |
| chr8 | 105271025 | *Hsf4* | DMR279 | NA | 0.0011655 | 0.0538632 |
| chr8 | 105271085 | *Hsf4* | DMR279 | NA | 0.004662 | 0.0372214 |
| chr8 | 105271093 | *Hsf4* | DMR279 | NA | 0.0011655 | 0.02695699 |
| chr10 | 66921261 | *Gm26576* | DMR11 | NA | 0.00815851 | 0.09247596 |
| chr10 | 66921282 | *Gm26576* | DMR11 | NA | 0.004662 | 0.07724434 |
| chr10 | 121150527 | *NA* | DMR21 | NA | 0.00815851 | 0.02768201 |
| chr11 | 72411971 | *NA* | DMR36 | NA | 0.00815851 | 0.0303354 |
| chr15 | 99199889 | *Spats2* | DMR108 | NA | 0.002331 | -0.0178079 |
| chr18 | 32430782 | *Bin1* | DMR142 | cg35563948 | 0.004662 | -0.0342284 |
| chr18 | 32430851 | *Bin1* | DMR142 | NA | 0.0011655 | -0.0325208 |
| chr18 | 32430889 | *Bin1* | DMR142 | NA | 0.004662 | -0.040947 |
| chr19 | 5783881 | *NA* | DMR148 | NA | 0.00815851 | 0.03218344 |

**Differentially methylated CpG sites analysis.** *Illumina.PROBEID*: probe name in the Illumina Infinium MouseMethylation Beadchip Array; Chrom and Position: probe location on mouse genome (assembly GRCm38/mm10); *DMR*: identity number of DMR; *Mann-Whitney*: Mann-Whitney nominal *p-value* obtained comparing CpG DNAm between Ts65Dn and wild-type euploid mice; *DeltaB:* difference between average Ts65Dn and wild-type euploid mice DNAm.
